# Regional radiomics similarity networks (R2SN) in the human brain: reproducibility, small-world and biological basis

**DOI:** 10.1101/2020.12.09.418509

**Authors:** Kun Zhao, Qiang Zheng, Tongtong Che, Dyrba Martin, Qiongling Li, Yanhui Ding, Yuanjie Zheng, Yong Liu, Shuyu Li

## Abstract

**Background:** Structural covariance network (SCN) has been applied successfully to structural magnetic resonance imaging (MRI) study. However, most SCNs were constructed by the unitary marker, which was insensitive for the different disease phases. The aim of this study is to devise a novel regional radiomics similarity network (R2SN) that could provide more comprehensive information in morphological network analysis.

**Methods:** Regional radiomics similarity network (R2SN) was constructed by computing the Pearson correlations between the radiomics features extracted from any pair of regions for each subject. We further assessed the small-world property of R2SN using the graph theory method, as well as the reproducibility in the different datasets and the reliability with test-retest analysis. The relationship between the R2SN and inter-regional co-expression of gene enriched was also explored, as well as the relationship with general intelligence.

**Results:** The R2SN can be replicated in different datasets, also regardless of using different feature subsets. The R2SN showed high reliability with the test-retest analysis (ICC>0.7). Besides, the small-word property (σ>2) and the high correlation with the gene expression (R=0.24, P<0.001) and the general intelligence was found by R2SN.

**Conclusion:** R2SN provides a novel, reliable, and biologically plausible method to understand human morphological covariance based on structural MRI.

**Impact Statement:** Imaging biomarkers are the cornerstone of modern radiology, and the development of valid biomarkers is crucial for optimizing individualized prediction in neurological disorders like AD. Thus, the development of the data mining method from neuroimaging is crucial for adding the biomarkers of disease. This study confirmed that R2SN provides a novel, robust and biologically plausible model and a new perspective for understanding the human brain, therefore. Thus, the R2SN has great promise in further study.

## 1. Introduction

Structural magnetic resonance imaging (sMRI) plays an important role in neuroscience where gray matter volume and cortical thickness etc. are the most popular brain morphological measures. However, most of the studies typically aanlyze single/several anatomical regions independently without taking associations among brain regions into consideration (Alexander-Bloch et al., 2013, Pichet Binette et al., 2020), regarding which, complex heterogeneous network patterns which characterize the brain by supporting information transformation are important for understanding complex brain cognitive function (Bullmore and Sporns, 2012, Alexander-Bloch et al., 2013), Specifically, structural covariance network (SCN), is often used to reconstruct brain structural network from sMRI based on the similarity of gray matter morphology (He et al., 2007, Tijms et al., 2012), and is commonly used to measure the association between inter-regions in the human brain with the morphological similarity He et al., 2007, Montembeault et al., 2012, Tijms et al., 2012, Montembeault et al., 2016, Zhang et al., 2017, Kong et al., 2018, Seidlitz et al., 2018, Spreng et al., 2019).

The SCNs typically consist of nodes and edges, representing the predefined brain regions and the statistical similarity between them based on the predefined morphological markers such as volume or thickness, etc. (Alexander-Bloch et al., 2013). Several methodologies have been introduced for the reconstruction of connectome maps based on sMRI either at the group level or the individual level. Specifically, SCN based on cortical thickness (He et al., 2007, He et al., 2008), or gray matter volume (Yao et al., 2010) was well studied at the group level, while the above-mentioned biomarkers were also developed to construct brain networks using single or multiple morphological features at the individual level (Tijms et al., 2012, Wee et al., 2013, Kong et al., 2014, Kim et al., 2016). All these methods have been used to investigate network alterations in brain-related diseases (Yao et al., 2010, Zheng et al., 2015, Bethlehem et al., 2017, Yu et al., 2018). Seidlitz and colleagues have proposed a morphometric similarity network that captures cortical cytoarchitecture and linked to individual cognitive performance Seidlitz et al., 2018). Despite the progress in constructing different brain networks, a well-validated and widely accessible model for mapping the brain network architecture of anatomically brain regions in an individual human brain is needed. Radiomics is a powerful method to extract more detailed information robustly (Li et al., 2019) including intensity and texture features from each brain region (Parmar et al., 2015, Gillies et al., 2016, Chaddad et al., 2018). However, there is a literature gap regarding the construction of the radiomics-based similarity network, as well as the associated attributes, which could be a feasible anatomical topological mapping of the individual brain.

In this study, the first aim is to develop a novel regional radiomics similarity networks (R2SN). Building on this foundation, the second aim is to explore the reproducibility, small-world property, and the biological basis of the R2SN, including the relationship between R2SN indices and the co-expression of gene network or fluid intelligence score. The results confirmed that R2SN provides a novel, robust and biologically plausible model and a new perspective for understanding the human brain, therefore, the R2SN has great promise in further study.

## 2. Materials and methods

### 2.1 Subjects

A total of 848 subjects from the Human Connectome Project (HCP, https://www.humanconnectome.org/) were included in our study where all subjects were cognitively normal controls (NC, age: 28.82±3.68, gender (M/F): 371/477, fluid intelligence: 16.53±4.86). HCP was established in 2009 with an overarching objective of studying human brain connectivity and its variability in healthy adults (Van Essen et al., 2012). All HCP subjects were collected by 3T MR scanners. In the HCP protocol, fluid intelligence was assessed using a form of Raven’s progressive matrices with 24 items. The detailed subjects’ information can be found in (https://www.humanconnectome.org/study/hcp-young-adult/document/1200-subjects-data-release), and also can be found in the previous study (Van Essen et al., 2012).

### 2.2 Data preprocessing and radiomics feature extraction

For each subject, the T1-weighted MRI image was aligned to Montreal Neurological Institute (MNI) space using a combined linear and non-linear registration (including N4 bias field correction) and resampled to 1 mm × 1 mm ×1 mm for further analysis (Xie et al., 2016) (Figure 1a). Then, forty-seven radiomics features in each brain region were extracted with each region defined upon AAL atlas (a total of 90 regions) (Tzourio-Mazoyer et al., 2002). The radiomics features consisted of 14 intensity features and 33 texture features (Figure 1b). All features were described in the study by Aerts and colleagues (Aerts et al., 2014) and implemented as in-house MATLAB scripts (https://github.com/YongLiulab/). The definitions and detailed descriptions of the radiomics features can be found in previous publications (Aerts et al., 2014, Feng et al., 2018, Zhao et al., 2020). Redundancy features, defined as features having a high correlation with other features (R>0.9), were removed before subsequent analysis. Therefore, a final feature matrix with 25 × 90 for each individual was obtained for further analysis (Figure 1c).

**Figure 1.**
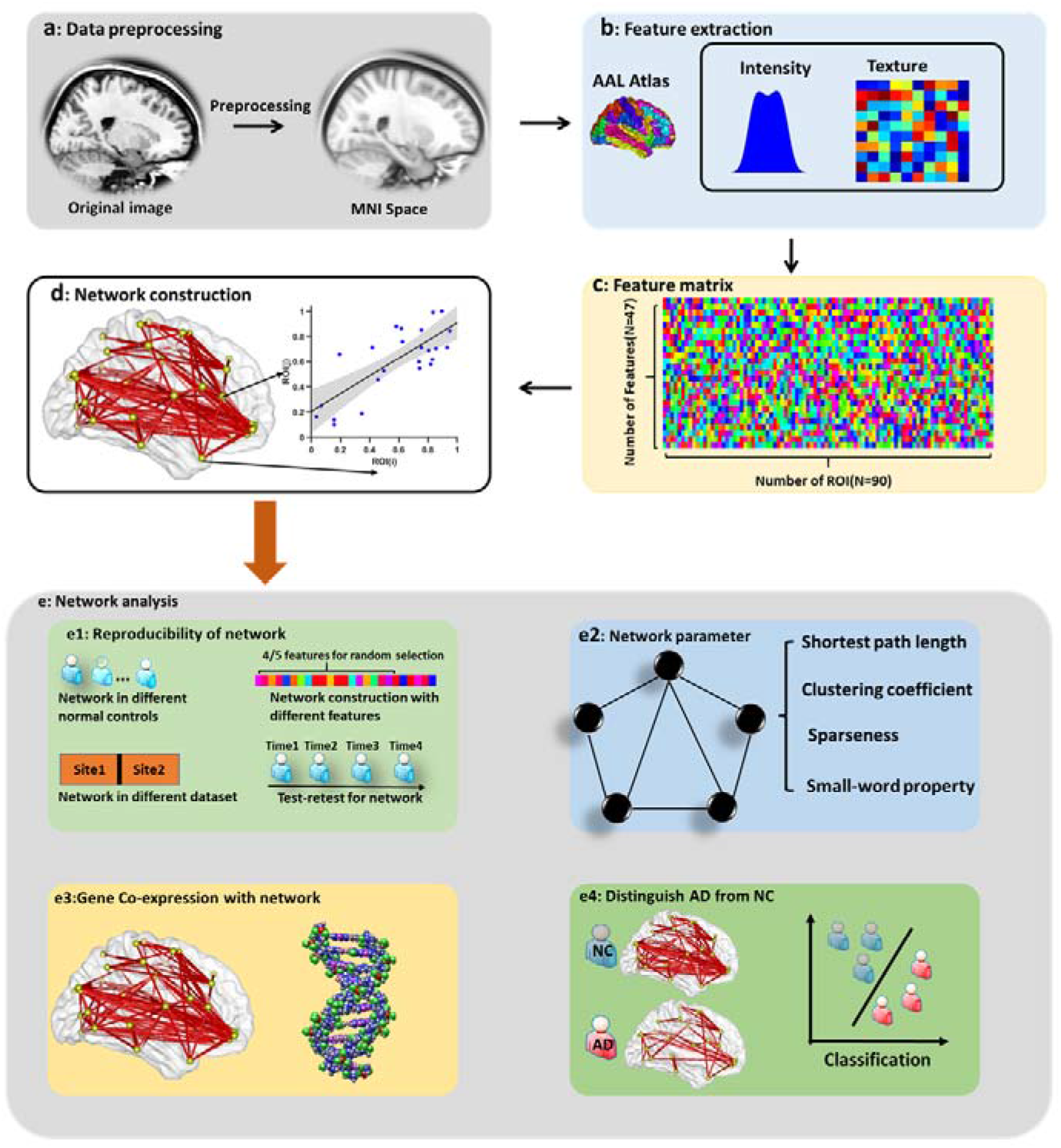
Schematic of the data analysis pipeline. (a) Data preprocessing. (b) The computation of radiomics features in each brain region. (c) Feature matrix of radiomics features. (d) Network construction with Pearson correlation. (e) The reproducibility of the network, (f) the network parameter of R2SN, (g) the correlation between R2SN and gene expression, and (h) the correlation between R2SN and fluid intelligence score.

### 2.3 R2SN construction

The R2SN was constructed by feature normalization, followed by a radiomics similarity matrix establishment. Specifically, the feature normalization was implemented by adopting a common min-max feature normalization scheme, while the radiomics similarity matrix establishment was performed by mapping the individual’s radiomics features into a radiomics similarity matrix of pairwise inter-regional Pearson correlations (Figure 1d). The mean value and the standard deviation (Std) of the R2SN were computed to estimate the fluctuation of R2SN in these young normal subjects (Figure 1e).

### 2.4 The topological structure of R2SN

To explore the topological structure of the brain characterized by R2SN, a variety of graph-theoretical network parameters were computed including shortest path length (L), clustering coefficient (C), and small-world property, after binarization of the R2SN using the threshold ranging from 0.5 to 0.75 (step size = 0.01). The detailed definitions of those parameters can be found in a previous study (Rubinov and Sporns, 2010) and computed by using the “Brant” (Xu et al., 2018) (Figure 1f). Briefly, the small-world index was defined as sigma (σ)=γ/λ, with the gamma being (γ) = C_R2SN_/C_random_, the lambda being (λ) = L_R2SN_/L_random_, C_random_ and L_random_ being the clustering coefficient and shortest path length of the random network (random sampling of edges to yield a matrix with the same number of connections) (Figure 1f).

### 2.5 Reproducibility of network

To test the reproducibility of R2SN among different datasets, the HCP dataset was randomly divided into two sub-datasets for 1000 iterations (424 subjects for each sub-dataset). The Pearson correlation coefficient of the two sub-datasets in each iteration was used to estimate the consistency of mR2SN, which was defined as the mean value of the R2SN in the specific dataset. At last, the distribution of the 1000 Pearson correlation coefficients was used to estimate the reproducibility of the R2SN.

Furthermore, to assess the reliability of the R2SN, a test-retest analysis was conducted using 21 subjects with 4 visits image (http://duke.edu/~morey005/ScanRescanData/). In this dataset, each subject was scanned on two different days, two scans were conducted 1h apart on day 1 (scans 1A and 1B) and two scans were conducted 1 h apart at a second session 7-9 days later (scans 2A and 2B) (Morey et al., 2010). The intra-class correlation coefficient [ICC, ICC = (BMS − WMS) / BMS] was used to estimate the reliability of each edge of the R2SN, where BMS is the between-subjects mean square, WMS is the within-subject mean square. The ICC has a value between 0 and 1, when ICC=0 means no reliability and ICC=1 means absolute reliability (Shrout. and Fleiss., 1979) (Figure 1e).

We also quantified the robustness of R2SN to methodological variations, including randomly reducing the number of radiomics features for analysis (i.e., only 20 radiomics features rather than all 25 features were involved as a predefined marker for network construction). The Pearson correlation coefficient was used to estimate the similarity of the mR2SN which was constructed with 20 radiomics features and 25 radiomics features. The distribution of the Pearson correlation coefficient of 1000 times simulation was performed to assess the robustness of R2SN to methodological variations (Figure 1e).

Besides, we explored whether the significant correlation can be obtained between mean connective strength and the size of each node (the size of each node was roughly estimated with the voxel number based on the AAL atlas) (Tzourio-Mazoyer et al., 2002).

### 2.6 The relationship between R2SN and Gene similarity network

To further explore the biological basis of the R2SN, we continue to compute the relationship between R2SN and the gene similarity network (GSN). The GSN was constructed with Allen atlas (http://human.brain-map.org/) (Zeng et al., 2012) and predefined genes (6 subjects). The nodes were defined as Allen atlas and mapped to the AAL brain regions based on MNI coordinate by using “abagen” (https://github.com/rmarkello/abagen) (Arnatkeviciute et al., 2019), and the edge was computed by the Pearson coefficient between the gene expression of any pair of regions. The Pearson’ correlation coefficient between the connectivities of the mR2SN and the connectivities of the mGSN (the mean value of the GSN) was calculated to assess the similarity between the R2SN and GSN (Figure 1g); furthermore, the Pearson’ correlation coefficient between the mean connect strengths of nodes of the mR2SN and mGSN to evaluate the similarity of these two networks.

### 2.7 Association between network properties of R2SN and cognitive difference

We have investigated the network properties, reproducibility, and biological basis of R2SN, and we assumed that the individual network properties could represent the individual’s differences in cognitive ability. In the HCP project, fluid intelligence (gF) scores were obtained as an index for measuring the subjects’ intrinsic cognitive ability. We focused on common general fluid intelligence (IQ) as the index to estimate the cognitive difference among young normal controls, similar to Seidlitz et al (Seidlitz et al., 2018). On this basis, we performed Pearson’s correlation analysis between fluid intelligence scores and connectivity strength and global efficiency of the individual’s network to explore the relationship between R2SN and fluid intelligence in the HCP dataset, and the global efficiency was computed with a “Brant” toolkit (Xu et al., 2018) (sparseness ranging from 0.2 to 0.8, step size=0.05) (Figure 1h).

## 3. Results

### 3.1 R2SN: edge properties

After redundancy removal, 25 radiomics features were reserved for each brain region, and the R2SN was constructed with those predefined features for each subject (a symmetrical matrix with a size of 90×90). The connective strength of the mR2SN was ranged from −0.56 to 0.99 in the HCP dataset (N=848), and the std value in each edge of the R2SN was significantly smaller than the mean value which was ranged from 0.01 to 0.36 but with more than 95% of them ranged from 0.01 to 0.20 (Figure 2a).

**Figure 2.**
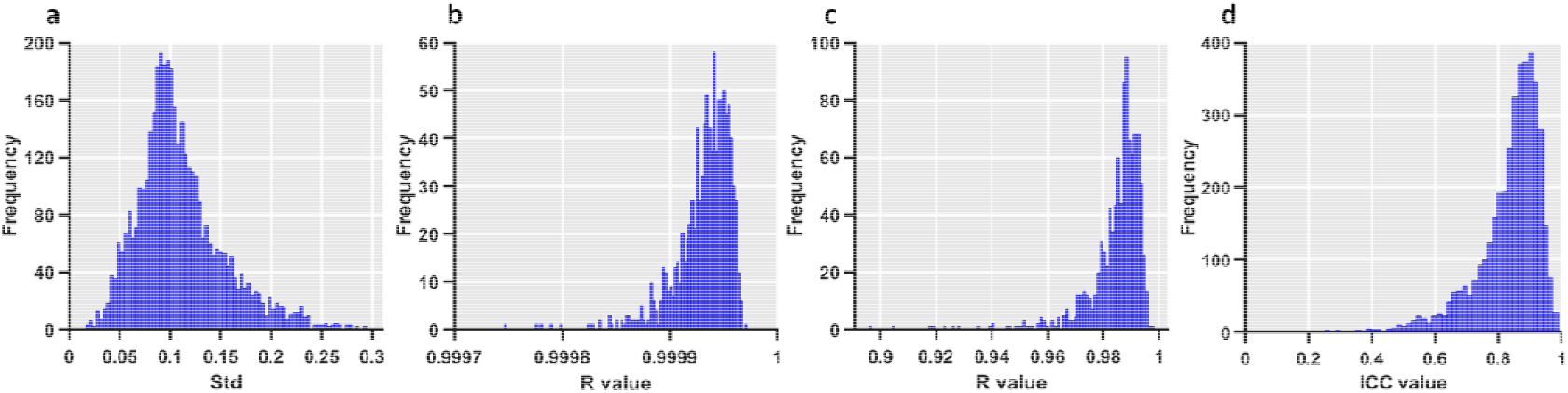
The stability and reliability of the R2SN. (a) The distribution of variance in the HCP dataset. (b) The distribution of the correlation coefficients about the R2SN network among each pair of two datasets (divide HCP dataset into 2 parts for 1000 times randomly repetitions). (c) The distribution of the correlation coefficients about the R2SN network with 20 features (random selection 20 features from 25 predefined features for 1000 times) and the R2SN which create with 25 features. (d) The ICC value of the R2SN with test-retest.

### 3.2 R2SN: network properties

The clustering coefficients, shortest path length, and the sparseness are shown in Figure 3a-c. Besides, the lambda (λ) showed the ratio of the shortest path length of R2SN and random network, and the gamma(γ) showed the ratio of the clustering coefficient of R2SN and random network. As a result, the value of λ was closed to 1 (Figure 3d) and the value of γ was significantly larger than 1 (Figure 3e), and the small world index sigma(σ) was also significantly larger than 1 by different thresholds of binarization (from 0.5 to 0.75, and the step size =0.01) (Figure 3f).

**Figure 3.**
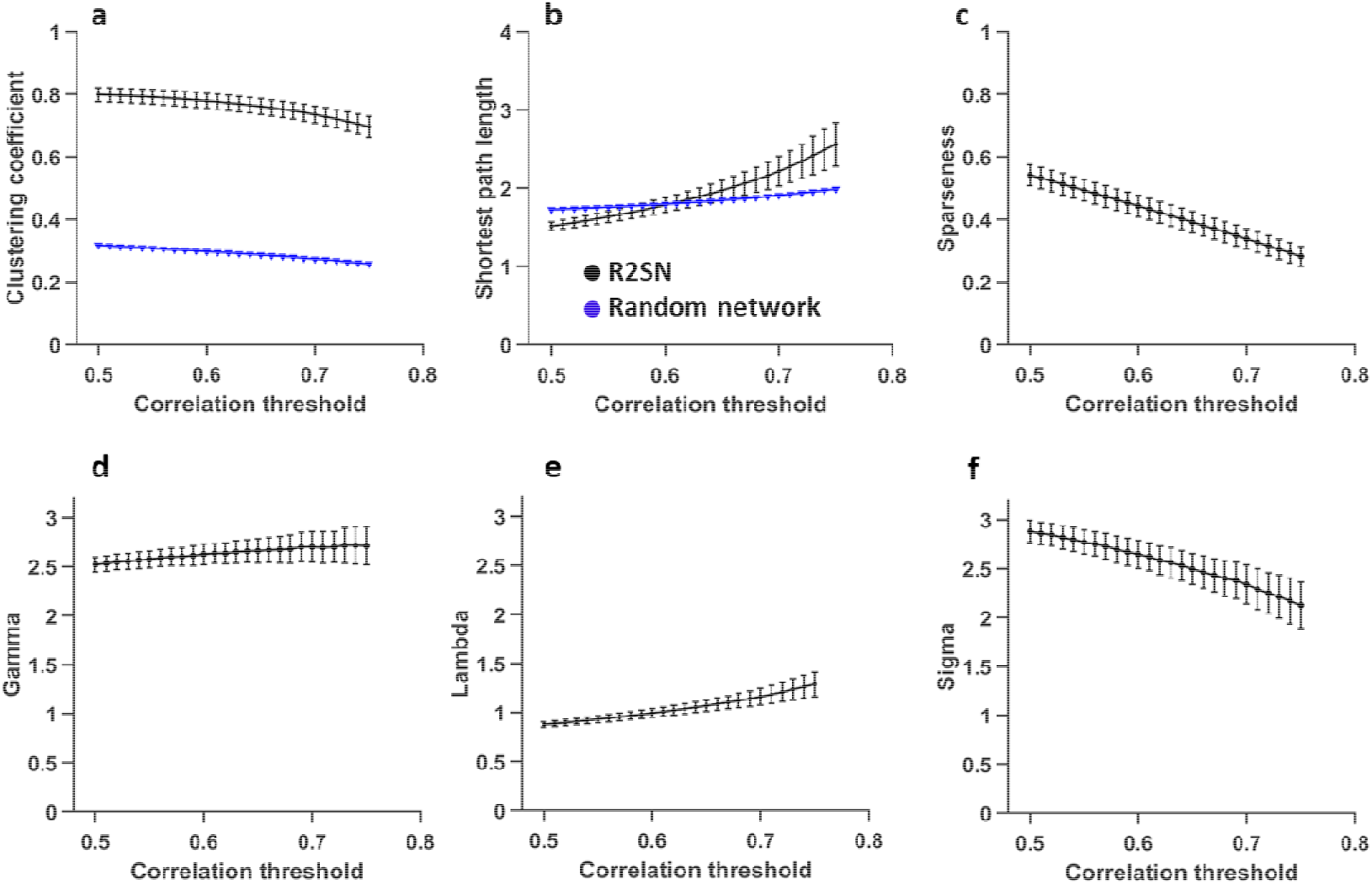
The network parameter about the R2SN with different correlation threshold (0.5-0.75 and step size=0.01), (a) the clustering coefficient, and the “black” means the R2SN, the “blue” means the random network, (b) shortest path length, (c) sparseness, and the small-world parameter, including (d) gamma value (the ratio of clustering coefficient between R2SN and random network), (e) lambda value and (the ratio of shortest path length between R2SN and random network) (f) sigma value (the ratio of gamma and lambda).

### 3.3 Reproducibility of R2SN

A serial of hypothesis testing was performed to further assess the reproducibility of the R2SN. Chief of all, the high consistency was found in any two mR2SNs which constructed by different dataset (1000 times randomly simulation) with the Pearson coefficient was ranged from 0.9992 to 0.9999 (Figure 2b). Besides, the high consistency also was obtained by mR2SNs which constructed with a different number of features (20 features for randomly selected and all features), and the R-value was ranged from 0.879 to 0.997 (Figure 2c). More importantly, the R2SN occurred high ICC value (ICC > 0.7) within more than 95% edge by test-retest analysis (Figure 2d). We do not find a significant correlation between the size of the node and mean connective strength (R =0.14, P =0.17).

### 3.4 The association between R2SN and Gene similarity network

For each subject from the Allen dataset, the GSN was constructed based on 15633 pre-defined genes (http://human.brain-map.org/). The mean connective strength of each node of mR2SN and mGSN are shown in Figure 4a (mR2SN), Figure 4b (mGSN), and Figure 4c. We also computed the similarity between mR2SN and mGSN (edge-based), a significant correlation can be found between two networks with R=0.27 (P<0.001), meaning that the brain region with high morphometric similarity also tended to have a high transcriptional similarity of the gene. We also computed the average of the off-diagonal elements of a row or column in the radiomics similarity matrix (node-based), the significant correlation also was found between mR2SN and mGSN with the mean connective strength of each node (R = 0.32, P =0.002).

**Figure 4.**
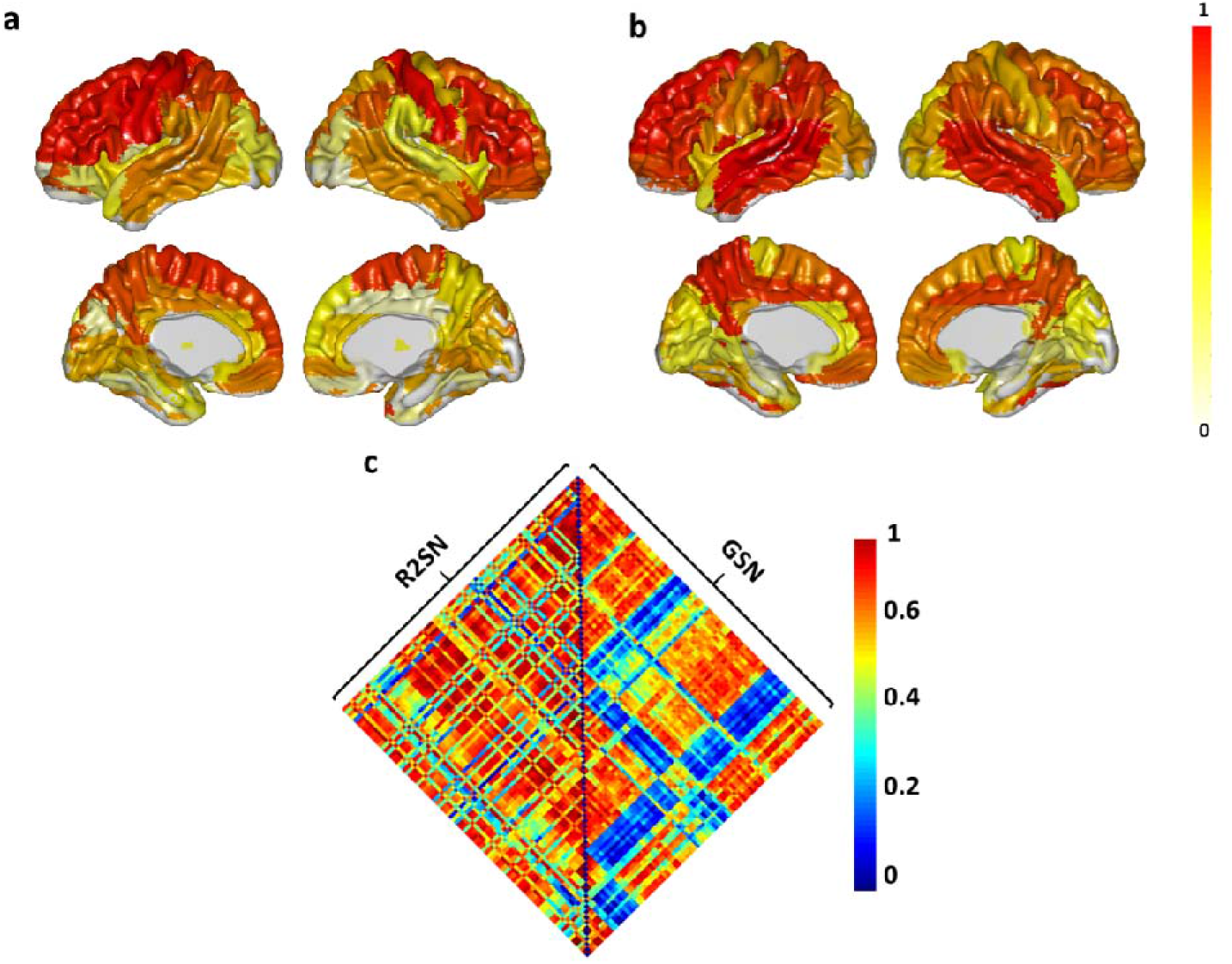
The correlation between R2SN and gene expression network. (a) The mean connective strength of R2SN, that was mapped to the surface area from the AAL template. (b) The mean connective strength of GSN, that was mapped to the surface area from the AAL template. the value of the color bar was normalized with the max-min method. (c) The hot-map for the R2SN and GSN. Some negative correlation was generated with GSN, and this phenomenon might be caused by the deletion of genes in some brain regions.

### 3.5 The association between R2SN and fluid intelligence

The fluid intelligence scores are range from 4 to 24 (https://www.humanconnectome.org/) for all subjects, we do not find a significant correlation between the mean strength connections/global efficiency of the overall R2SN network and fluid intelligence scores (P>0.05). The Pearson correlation showed that about 14% of connections have a significant correlation with fluid intelligence score (P<0.05, uncorrected), especially in the Precentral (left and right), Olfactory (left), Superior orbital gyrus (right), Rolandic operculum (right), Medial and lateral cingulate gyrus (left), Superior orbital frontal gyrus (right), Pallidum (right) and middle temporal gyrus (right) with P<0.05 (Bonferroni-corrected, N=4005) (Figure 5a). Briefly, the stronger the functional connections, the higher correlation with fluid intelligence score occurred. Besides, we also explored the relationship between fluid intelligence score and the global efficiency of each brain region, as figure 5b showed, the global efficiency of 14 nodes was significantly correlated with fluid intelligence (P<0.05) (when sparseness=0.55), especially in the Superior orbital gyrus (right), Rolandic operculum (right), Lingual (left) and Caudate (right) with P<0.005 (Figure 5b).

**Figure 5.**
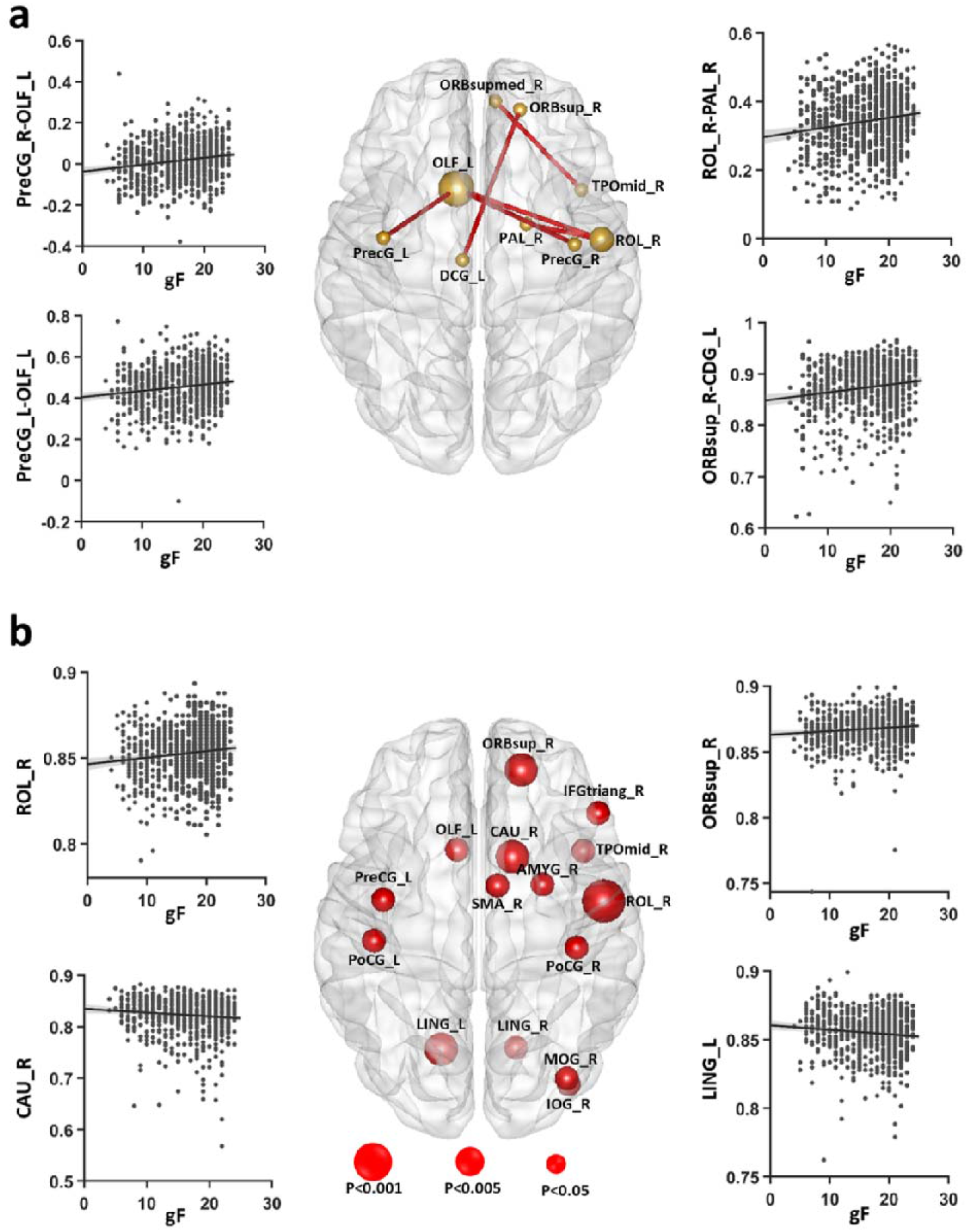
The correlation between R2SN and fluid intelligence score. (a) the connectivity which showed significant correlation with the individuals’ fluid intelligence score (P<0.05, Bofforni corrected), (b) the brain regions in which the global efficiency showed significant association with fluid intelligence score (P<0.05), the size of the red nodes reflected the P-value in 3 levels (<0.001, <0.005 or <0.05), and the scatter diagram for the node with P<0.05/4005 are provided as an example figure.

## 4. Discussion

The present study provides a pipeline for transforming radiomics feature maps into a radiomics similarity matrix of pairwise inter-regional at the individual’s level, which still represents a gap according to the literature. The systematic results confirmed that R2SN provides a novel, robust and biologically plausible model for understanding the human brain.

Radiomics is a powerful method to extract more detailed information from brain image, which includes intensity, and texture features, which could improve SCN in representing the morphology of the brain as a network with the relativeness of pair of regions (Parmar et al., 2015, Gillies et al., 2016, Chaddad et al., 2018). Texture features can quantify the variations in intensity or patterns, including those features that are imperceptible to the human visual system (Aerts et al., 2014). Numerous studies emphasized the importance of radiomics (Gillies et al., 2016), and also took it as a bridge between imaging and personalized medicine (Lambin et al., 2017). More importantly, radiomics also improve the power of the precision of diagnosis, treatment, and prognosis of the tumor (Aerts et al., 2014, Coroller et al., 2015, Huang et al., 2018), and diagnosis of AD (Sorensen et al., 2016, Sorensen et al., 2017, Feng et al., 2018, Zhao et al., 2020). Hereby, the radiomics is a method that extracts a large number of features of the brain region with high reproducibility (Li et al., 2019) that provide a better measure to facilitate better characteristics of the brain region.

Representing the morphology of the brain as a network has the advantage that the structure of the brain can be described statistically with tools from graph theory (Tijms et al., 2012). Like most of the brain networks (e.g. functional brain network, diffusion-weighted imaging (DWI), and SCN) (He et al., 2007, Tijms et al., 2012, Heinze et al., 2015, Seidlitz et al., 2018), the R2SN also had a complex topology. In the R2SN, the high-degree nodes were located in the frontal lobe, parietal lobe, and occipital lobe, and low-degree nodes were found in the temporal lobe and subcortical. The nodes which occurred higher connections might have more cooperation with other brain regions and the node which occurred lower connection might have a more specific function in the brain (Seidlitz et al., 2018). The different connective patterns between the GSN and R2SN might be caused by the difference of microstructure in the brain like the histological classification of cortical areas (Solari and Stoner, 2011, Seidlitz et al., 2018).

Notably, R2SN can be replicated in different datasets, demonstrated by a subsampling strategy from the same dataset for 1000 iterations. The R2SN showed high intra-class class correlation coefficients (ICC) (a prominent statistic to measure test-retest reliability) in different visit images, indicating the robustness of the radiomics similarity connectivity under the consensus that the reproducibility was the most important property for a novel method in MRI analysis (Baker, 2016). Therefore, the high consistency of the R2SN which was constructed with different features also confirmed the methodological variations (Seidlitz et al., 2018). Those properties of the R2SN was to support the firm foundation for the credibility of the results.

Imaging biomarkers are taken as the cornerstone of the radiology community and imaging genetics has established heritable phenotypes for quantitative genetics of brain phenotypes. As expected, a high correlation was found between gene expression network and R2SN (Zeng et al., 2012), meaning that cortical areas with high morphometric similarity also tended to have high transcriptional similarity (Seidlitz et al., 2018). The structure of the brain region was controlled by gene expression (Liu et al., 2004, Warden and Mayfield, 2017), and the variations in intensity or pattern of the brain region can be reflected by the radiomics feature (Wang et al., 2019), these results further speculated the gene expression can be reflected with the R2SN. In brief, R2SN has a genetic basis, and those findings provided the potential possibility to estimate the risk of the gene by using R2SN and also provided certain evidence for genetic disease research, such as Alzheimer’s disease (Zhao et al., 2020).

It is vital to evaluate the association between the individual network architecture and the cognitive ability or psychological functions of the brain (Li et al., 2009, van den Heuvel et al., 2009, Seidlitz et al., 2018). High-degree hub nodes and the global efficiency of the connectome to be preferentially affected by clinical brain disorders associated with cognitive impairment, the relationship between high-degree nodes and cognitive status might be obtained a higher performance in classification, prediction, and so on (Crossley et al., 2014). As we all know that fluid intelligence is a common measure to estimate the cognitive ability among normal controls and to identify the cognitive difference among individuals, the higher fluid intelligence scores indicated correspond to more efficient information transfer in the brain (Li et al., 2009). The fluid intelligence score refers to the ability to solve abstract problems that do not depend on acquired knowledge and changes with age (Gray et al., 2003, Cole et al., 2012, Kievit et al., 2016). A set of brain regions frontoparietal have been reported associated with fluid intelligence in brain imaging (Duncan et al., 2000, Cole et al., 2012, Woolgar et al., 2018). Convergence evidence indicated that the lingual gyrus, caudate nucleus, rolandic operculum, and the frontal lobe which play an important role in an individual’s cognitive ability (Preusse et al., 2011, Rhein et al., 2014, Santarnecchi et al., 2015, Zhang et al., 2016). The significant correlation between fluid intelligence and efficiency of the hub-regions in the R2SN also indicates an individual’s cognitive ability linked to the brain network architecture (Li et al., 2009, van den Heuvel et al., 2009).

### Limitation and caveats

This study has some limitations. First, we just study the R2SN based on AAL Atlas, the network properties and its basis need to be further validated by other fine Brain Atlas. Secondly, a unified framework for interpreting these measures and their alterations in different brain diseases is needed. Thirdly, more samples from different independent scanners, more cognition measures may improve the statistical power of the analysis, allowing scientists to explore the neural mechanisms of R2SN in the future.

## 5. Conclusion

R2SN is a network with high stability, reproducibility, and a biological basis, which might improve a novel method and shed new light on future MRI studies. We assume that R2SN could provide a powerful technology platform for measuring the anatomical connectome in vivo and be applied to the diagnosis of a variety of diseases in the future.

## 6. Acknowledgments

This work was partially supported by the National Key Research and Development Program of China (grant no. 2017YFB1002502), the National Natural Science Foundation of China (grant nos. 81972160, 81622025, 81871438, 61802330, 81871508, 61773246), Beijing Natural Science Funds for Distinguished Young Scholar (JQ200036), and Open Project Program of the National Laboratory of Pattern Recognition (NLPR) (No. 201900021). Taishan Scholar Program of Shandong Province of China (No. TSHW201502038) and Major Program of Shandong Province Natural Science Foundation (ZR2019ZD04, No. ZR2018ZB0419).

## Author contribution

Kun Zhao analyzed the data and performed the measurements; Kun Zhao, Yong Liu and Shuyu Li were majorly responsible for preparing the manuscript. Qiang Zheng, Tongtong Che, Dyrba Martin, Qiongling Li, Yanhui Ding, Yuanjie Zheng, Shuyu Li, and Yong Liu revised the paper, Shuyu Li and Yong Liu supervised the project.

